# Draft Genome Sequences of Seven Strains of *Dickeya dadantii*, a Quick Decline-causing Pathogen in Fruit Trees, Isolated from Japan

**DOI:** 10.1101/2020.05.24.113969

**Authors:** Takashi Fujikawa, Hiroe Hatomi, Nobuyoshi Ota

## Abstract

Plant pathogenic bacterium *Dickeya dadantii* causes quick decline in fruit trees (apple, Japanese pear, and peach). In this study, we report on the draft genome sequences of seven strains of *D. dadantii* isolated from fruit trees with typical quick decline symptoms in Japan.

---

*Dickeya dadantii* is a plant pathogenic bacterium that causes soft rot disease in various plants (1). In addition to being a soft-rot pathogen, it causes quick decline (QD) in fruit trees, such as apple, peach, and Japanese pear trees (2, 3). The symptoms of QD include red-brown sap leakage from the trunk and/or branches, softening bark, defoliation, leaf and shoot necrosis, and dieback. However, the lifecycle of *D. dadantii* is not yet well understood. The genetic characteristics of this bacterium will need to be defined to develop effective measures for the control of QD. To this end, we performed whole-genome sequencing (WGS) of seven *D. dadantii* strains isolated from fruit trees with symptoms of QD.

The strains sequenced in this study were obtained from the Institute of Fruit Tree and Tea Science (NARO) (Table 1), isolated from apple, peach, and Japanese pear trees, and identified as *D. dadantii* according to our previous studies (2, 3). The strains were cultivated in YP broth (2) at 27°C for 1 day with agitation at 140 rpm. Then, 1-ml aliquots of each culture were used for DNA extraction with a DNeasy mini kit (Qiagen, Hilden, Germany). Their genome sequences were determined using WGS, as previously reported (4, 5, 6, 7). Briefly, the genomic DNA was sequenced using an Ion PGM sequencer with an Ion PGM Hi-Q View OT2 kit, an Ion PGM Hi-Q View Sequencing kit, and a 318 Chip kit v2 (Thermo Fisher Scientific, Waltham, MA, USA), according to the manufacturer’s instructions. Default parameters were used unless specified otherwise. The sequence reads were quality controlled (quality score <20) and the adapter sequences were removed using the CLC Genomics Workbench (version 10, except for Kunimi3-1 (ver. 12), and BI1-1 (ver. 20)). Using the resulting reads, contigs (filtered with a size >500 bp) were assembled *de novo* using the CLC Genomics Workbench. The draft genomes were annotated using the NCBI Prokaryote Genome Annotation Pipeline (PGAP).

**Table 1.** Genome data and accession numbers of seven *Dickeya dadantii*.

|  | Strain information |  | Genome information |  |  |  |  | PGAP<sup>a</sup> annotation |  |  | Reads information |  |  |  |
| --- | --- | --- | --- | --- | --- | --- | --- | --- | --- | --- | --- | --- | --- | --- |
| Strain | Isolation host | Isolation area | GenBank accession no. | Genome size (bp) | G+C content (mol% ) | No. of contigs | $N_{50}$ | Total no. of genes | rRNAs (5S, 16S, 23S) | tRNAs | SRA<sup>b</sup> accession no. | No. of reads | Average length (bp) | Genome coverage (×) |
| BI1-1 | Apple | Japan: Iwate | JABEOZ000000000 | 4,663,807 | 56.2 | 271 | 39,637 | 4,311 | 1, 2, 1 | 46 | SRR11696188 | 673,373 | 229.2 | 33.1 |
| BI3-1 | Apple | Japan: Iwate | PHRA000000000 | 4,771,676 | 56.3 | 92 | 151,183 | 4,331 | 4, 1, 1 | 61 | SRR11692751 | 5,046,907 | 298.9 | 316.1 |
| Aka1-1 | Peach | Japan: Fukushima | JABEPA000000000 | 4,714,768 | 56.4 | 268 | 43,532 | 4,429 | 5, 1, 1 | 57 | SRR11696187 | 592,688 | 258.8 | 32.5 |
| Yana2-2 | Peach | Japan: Fukushima | JABEPB000000000 | 4,825,596 | 56.4 | 204 | 61,681 | 4,432 | 3, 2, 1 | 48 | SRR11696186 | 1,510,292 | 255.4 | 79.9 |
| Kunimi-3 | Peach | Japan: Fukushima | SMHE000000000 | 4,869,298 | 56.5 | 141 | 96,135 | 4,446 | 2, 1, 2 | 62 | SRR11730645 | 1,941,545 | 287.6 | 114.7 |
| Kousui1-1 | Japanese pear | Japan: Saga | JABEPC000000000 | 4,881,710 | 56.1 | 193 | 68,290 | 4,488 | 5, 1, 3 | 53 | SRR11696185 | 1,308,248 | 263.2 | 70.5 |
| Housui2-1 | Japanese pear | Japan: Saga | JABEPD000000000 | 4,882,493 | 56.2 | 197 | 56,134 | 4,481 | 4, 1, 6 | 49 | SRR11696184 | 1,746,612 | 258.4 | 92.4 |

<sup>a</sup>NCBI Pipeline Genome Annotation Pipeline
<sup>b</sup>short read archive

The WGS analysis indicated that the genome size of the seven strains was 4.7-4.9 Mbp with a G+C content of 56.1-56.4% (Table 1). The genome information of this species was previously published in NCBI as 4.7-5.0 Mbp with a G+C content of 56.3-56.5% (e.g. strains 3937 and DSM 18020; GenBank accession no. CP002038 and CP023467, respectively), which supports the results of our WGS analysis. Moreover, PGAP identified 4,311-4,488 genes and multiple rRNA and tRNA genes in these genomes (Table 1). This information can be used to compare the genomes and gene expression patterns of different strains or species. Therefore, the results of our WGS analysis may help to elucidate the characteristics of *D. dadantii*, a bacterium related to the virulence of QD.

## Data availability

All WGS projects in this study have been deposited at DDBJ/ENA/GenBank. The corresponding read data are available from the Sequence Read Archive (SRA) with the accession numbers provided in Table 1.

## Acknowledgments

We would like to express our gratitude to Ms. M. Taguchi and Ms. A. Sasaki for supporting this work and to the members of IFTS-NARO for their helpful discussions. We would also like to thank Editage (www.editage.jp) for English language editing. This work received no specific grant from any funding agency in the public, commercial, or not-for-profit sectors.

